# Permutation tests for comparative data

**DOI:** 10.1101/2020.10.24.325472

**Authors:** James G. Saulsbury

## Abstract

The analysis of patterns in comparative data has come to be dominated by least-squares regression, mainly as implemented in phylogenetic generalized least-squares (PGLS). This approach has two main drawbacks: it makes relatively restrictive assumptions about distributions and can only address questions about the conditional mean of one variable as a function of other variables. Here I introduce two new non-parametric constructs for the analysis of a broader range of comparative questions: phylogenetic permutation tests, based on cyclic permutations and permutations conserving phylogenetic signal. The cyclic permutation test, an extension of the restricted permutation test that performs exchanges by rotating nodes on the phylogeny, performs well within and outside the bounds where PGLS is applicable but can only be used for balanced trees. The signal-based permutation test has identical statistical properties and works with all trees. The statistical performance of these tests compares favorably with independent contrasts and surpasses that of a previously developed permutation test that exchanges closely related pairs of observations more frequently. Three case studies illustrate the use of phylogenetic permutations for quantile regression with non-normal and heteroscedastic data, testing hypotheses about morphospace occupation, and comparative problems in which the data points are not tips in the phylogeny.

## Introduction

For a biologist interested in the role of natural selection in evolution, questions about relative trait values are easier to address than questions about absolute trait values. For example, “do bears from colder climates have longer fur” is far more analytically tractable than “is long hair an adaptation for cold climates,” even if the latter is the original question of interest (Sober and Orzack 2003). Comparative or cross-species data are a fruitful source of insights into how natural selection works in populations, and also into broad-scale phenomena that are interesting in themselves, but their analysis is non-trivial. Comparative data often carry a detectable signal of the phylogenies on which they evolved, and covariation between the trait values of close relatives can cause serious problems for a statistical analysis, most conspicuously in the form of inflated false positive rates (Felsenstein 1985). The dominant paradigm for the past several decades of comparative research was established by Felsenstein (1985), who showed that the independent values in a comparative analysis are not the trait states at the tips of a phylogeny but their divergences (or contrasts) at phylogenetic splits. Unlike the raw trait values, these “phylogenetically independent contrasts” (PICs) can be safely analyzed with least-squares regression. Phylogenetic generalized least squares (PGLS; Grafen 1989) was developed as a more general comparative framework that can accommodate non-linear relationships via link functions (for example, phylogenetic logistic regression; Ives and Garland 2010), trees with polytomies, and a variety of evolutionary models. PGLS is a kind of generalized least squares regression that uses a phylogeny as the variance-covariance matrix, and it returns identical results as the PIC approach in its simplest form.

PICs/PGLS have enjoyed immense success as a framework for understanding relationships among traits in comparative data while accounting for phylogenetic autocorrelation (Symonds and Blomberg 2014), but they have two chief limitations. First, as regression tests they are assumption-rich: their reliability depends on, among other things, the residuals being normally distributed and homoscedastic (equal variance across the values of the predictors) (Mundry 2014). The other limitation is that least-squares regression is a rather specific analytical framework: questions about the relationship between one or more variables and the conditional mean of another variable occupy only a small corner of the universe of biologically interesting comparative problems. This has pernicious implications for the use of phylogenetic regression as the “go-to” method among comparative biologists. In the last section of this paper I highlight three examples of comparative problems that are off-limits to PGLS: quantile regression, morphospace occupation, and ecogeographic rules. PICs/PGLS is quite powerful within rather circumscribed bounds (Orzack and Sober 2001), but methods that promote the creative exploration of questions and datasets outside those bounds can only improve comparative biology. Here I develop phylogenetically informed permutation tests, validate them with toy scenarios, and illustrate their use with empirical case studies. The first of these tests generates a set of nulls using cyclic permutations and is a conceptually straightforward extension of an existing test, but it can only be used with balanced trees. The second conserves the phylogenetic signal in the data, has identical statistical properties to the first, and can be used with any phylogeny.

### Phylogenetic permutation tests

Permutation testing is widespread for some questions in evolutionary biology – for example, in tests of phylogenetic signal (Blomberg et al. 2003) – but strangely has not permeated to comparative testing (with one important exception, discussed below). The gist of a permutation test is to take as a null distribution the set of test statistics associated with every unique rearrangement (or permutation) of the data and to compare the empirical test statistic with this distribution (Good 2000). In each of these permutations the “labels” on at least one variable in the dataset are randomly rearranged, breaking the empirical association between variables without changing their distributions. The proportion of permutations in which the test statistic is at least as extreme as the empirical one is taken as the probability of obtaining the results under the null hypothesis: the p-value (Perezgonzalez 2015). For example, the subject in Fisher’s famous “Lady Tasting Tea” experiment correctly guessed the method of preparation for 8 cups of tea, and Fisher used permutations of the order of guesses to determine that guessing randomly would have achieved this result with a low probability of p = 1/70. This example was simple enough for every possible permutation to be enumerated analytically, but for bigger datasets this can be computationally infeasible, so in practice the distribution is typically estimated by permuting randomly many times (Good 2000). Many “flavors” of permutation test have been developed, differing mainly in the null distribution they generate (Anderson 2001). The primary virtue of the permutation test is its elegance: unlike parametric tests, it does not rely on theoretical probability distributions (the population of interest is the empirical one), and it can be used with a broader range of test statistics (Good 2000).

Despite its strengths, the ordinary permutation test cannot be applied to data that evolved on phylogenies. The test is, in a sense, distribution-free, but not assumption-free: permutation tests assume among other things that the observations being shuffled are *exchangeable*, meaning that rearrangements of those observations have the same joint probability distribution (Anderson 2001). This is quite close to the assumption in least-squares regression that variables are independent and identically distributed, and these assumptions are violated by the complex covariance structure of comparative data. In other words, because the traits of closely related taxa tend to covary due to their shared evolutionary history, comparative data are not exchangeable. However, a modified test that uses phylogenetic information to preserve exchangeability can be used for sound non-parametric hypothesis-testing with comparative data.

#### Lapointe-Garland phylogenetic permutations

Lapointe and Garland (2001) proposed a permutation test for comparative data in which pairs of values at the tips are exchanged with probability proportional to their phylogenetic proximity, such that the most probable exchange is between a trait value and itself (Fig. 1A). This approach uses a relatedness matrix which can be “flattened out” using a parameter k; for values of k higher than one the test approaches an ordinary permutation test. This was the first and apparently the only previous attempt at developing a comparative permutation test.

**Figure 1.**
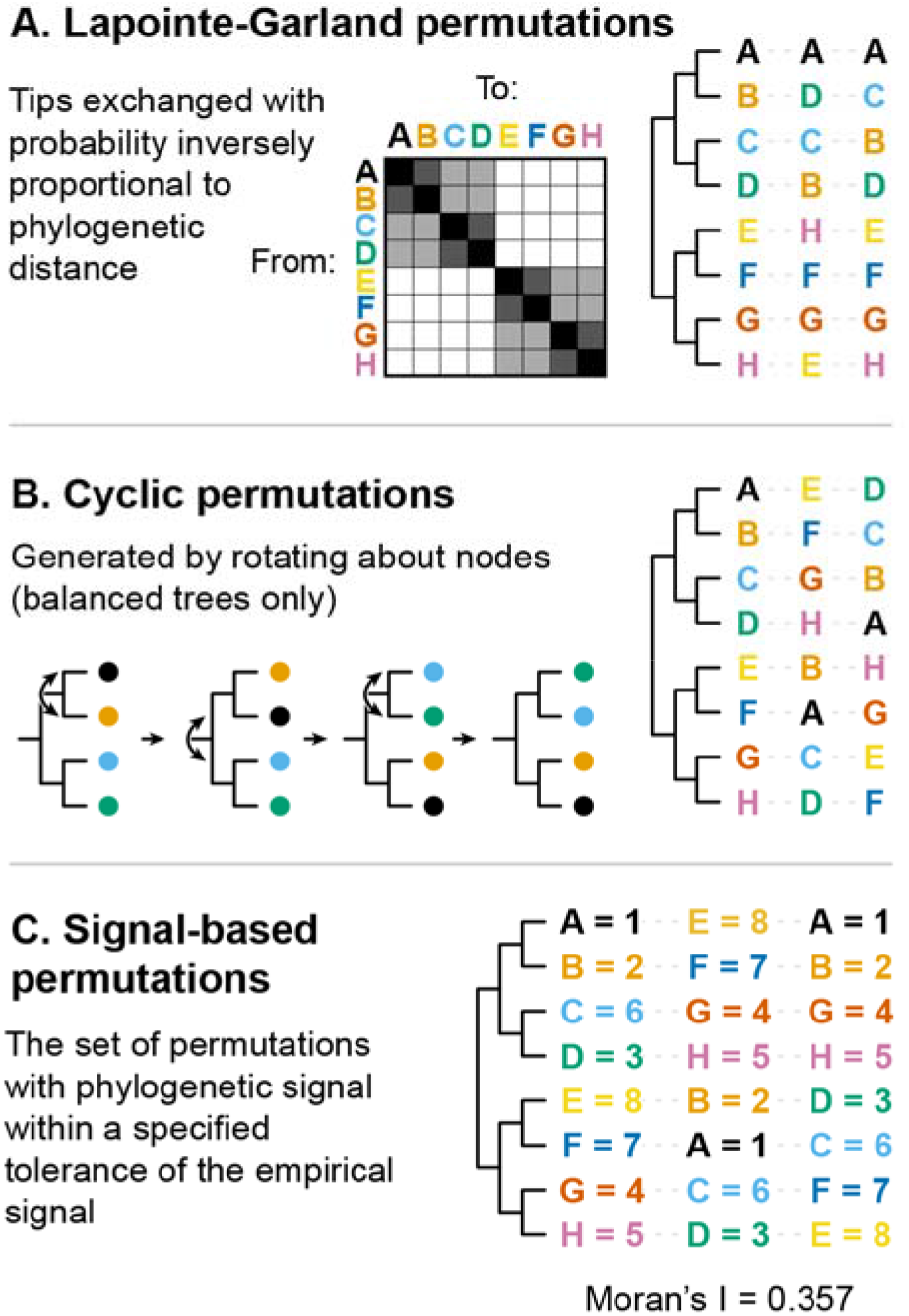
The three kinds of phylogenetic permutations discussed in this paper, with examples of each kind on a balanced, rooted tree of 8 taxa A-H. In a test, each kind of permutation is applied to at least one variable in the dataset, and a population of many such permuted datasets is compared with the empirical arrangement. **1A.** The phylogenetic permutation approach developed by Lapointe and Garland (2001). Note that many trait values do not change position across permutations because the highest probability of exchange is between a trait and itself. **1B.** Cyclic permutations: the set of permutations that can be generated by rotating about nodes in the tree (double-sided arrows). **1C.** Signal-based permutations. These are more inclusive than cyclic permutations: they include all possible cyclic permutations because the latter always conserves phylogenetic signal, but also non-cyclic permutations that retain the same or nearly the same signal (rightmost rearrangement). This is the only test in which the set of permutations depends on the values at the tips.

Although the test is less vulnerable than the ordinary permutation test to phylogenetically induced false positives, it has several undesirable properties. One of these is the high rate of exchanges between an observation and itself (Fig. 1A), or auto-exchanges, which results in a set of permutations that is tightly constrained around the empirical statistic (Appendix 1). This should reduce statistical power because the permutations look so much like the empirical arrangement. In this respect the Lapointe-Garland (LG) test also strays from one of the essential features of permutation tests: enumerating each unique rearrangement of the data. The high rate of auto-exchanges has the effect of up-weighting some possible rearrangements over others; it is not clear why it would be desirable to give more weight to rearrangements that look more like the empirical one. The rate of auto-exchanges can be dampened by increasing the value of k, which makes the exchange matrix flatter, but that defeats the point of incorporating phylogeny. Moreover, there is no principled way to choose a value of k above one, nor is there a clear use-case for permutations that are only partially informed by phylogeny. Another theoretical problem with LG permutations is that they do not conserve phylogenetic signal, the key feature that makes interpreting comparative datasets difficult. For the leftmost tree in Fig. 1C, the phylogenetic signal of LG permutations varies by a factor of 2.3.

Simulations show these features of the LG approach have consequences for its statistical performance. For instance, one desirable feature of a significance test is that the rate of false positives should be exactly equal to the significance level: thus, 5% of cases in which the null hypothesis is true should have p < 0.05. I tested the false positive rate of the LG permutation test by simulating uncorrelated Brownian Motion evolution of two continuous traits on a rooted 8-taxon tree with two polytomies of four tips each. I computed the absolute correlation coefficient between the two simulated traits and tested it with an LG permutation test (500 permutations) for each of 1000 pairs of simulated traits (Fig. 2). I ran the same test with independent contrasts, after making the tree amenable to PICs by making the two 4-taxon polytomies in the tree into pectinate subtrees with added branches having length 0. Unlike independent contrasts, LG permutation tests yielded non-uniformly distributed p-values, returning intermediate values most frequently (Kolmogorov-Smirnov test against a uniform distribution, p = 0.011). In other words, the p-values from this test do not tell the user what they are supposed to: the probability of observing a statistic at least as extreme as the empirical one under the null hypothesis. I also evaluated the false negative rate with the same procedure, except the two simulated traits were correlated with an evolutionary covariance of 0.75. LG permutation tests return false negatives at higher rates than independent contrasts: p < 0.05 for 469 of 1000 simulations of truly correlated evolution, compared with 564. The LG phylogenetic permutation test is a valuable and interesting non-parametric approach to comparative data, but, motivated by the conceptual and statistical problems outlined here, I develop two new phylogenetically informed permutation tests. The cyclic permutation test is a conceptually straightforward extension of an existing class of permutation tests that can only be used with balanced trees; the signal-based permutation test has identical statistical properties to the first and can be used with any phylogeny.

**Figure 2.**
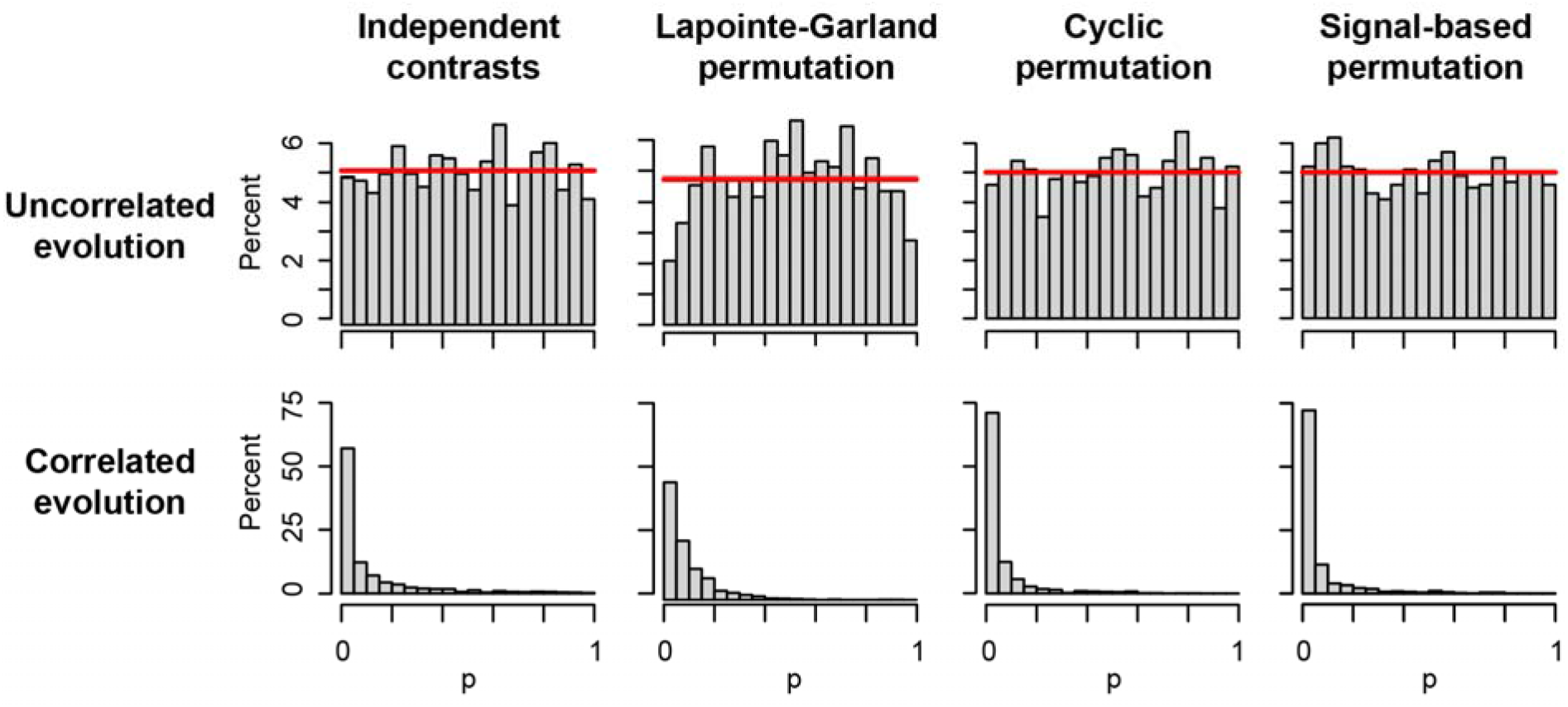
p-values for three phylogenetic permutation tests of correlation and one test with independent contrasts, applied to uncorrelated evolution of X and Y (above) and correlated evolution with an evolutionary covariation of 0.75 (below). Traits simulated on a rooted 8-taxon tree containing two polytomies with 4 taxa each, all branch lengths equal. p-values should ideally be uniformly distributed for uncorrelated evolution and as low as possible for correlated evolution. Red horizontal line indicates a uniform distribution; only Lapointe-Garland permutations differ significantly from this distribution. All bins have width 0.05.

#### Cyclic permutations

An elegant solution to the problem of relatedness in comparative data can be found in the restricted permutation test, in which rearrangements are restricted to only occur between exchangeable data points or sets of data points (Anderson 2001). As a non-phylogenetic example, an investigator testing the significance of a correlation between environmental variables sampled in different regions might consider permuting only within regions and not across them, especially if those variables were spatially autocorrelated by region. The resulting restricted permutations would retain the same kind of spatial autocorrelation as the empirical data. In a comparative dataset, the exchangeable units are not the values at the tips but the descendants of each node in the tree. This is similar to Felsenstein’s (1985) observation that contrasts at nodes rather than tip values are independent of one another, and similar also to the “radiation principle” that motivates Grafen’s (1989) PGLS. A set of phylogenetically informed permutations can therefore be generated with cyclic permutations of the values at the tips; that is, by randomly rotating the descendants of each internal node for at least one variable in the dataset (Fig. 1B). A statistic calculated for the set of these permuted datasets can be compared with the empirical one in what is here called a cyclic permutation test. Because the units being permuted can either be tip values (for the shallowest internal nodes) or sets of tip values (for deeper nodes), this is a form of hierarchical restricted permutation.

The cyclic permutation test performs at least as well as independent contrasts and lacks the statistical issues of the LG permutation test. In the test for false positives (Fig. 2), the set of p-values is indistinguishable from a uniform distribution (Kolmogorov-Smirnov test, p = 0.370), which is ideal. In the test for false negatives with cyclic permutations, p was below 0.05 for 702/1000 simulations, corresponding to a false negative rate of around 30% (Fig. 2). Interestingly, this is a better rate than what independent contrasts recovered (p below 0.05 for 564/1000 simulations). This indicates that the cyclic permutation test and PICs have at least comparable statistical power, even though the former is a non-parametric test.

Cyclic permutations will change an unbalanced tree’s two-dimensional projection, so the cyclic permutation test can only be used with a topologically balanced tree. If a trait is permuted cyclically on an unbalanced tree, it will no longer share the same evolutionary history as other traits in the dataset, and it defeats the point of the test – namely, to ask what kinds of patterns can result from the independent evolution of different traits on the same phylogeny. Because of the restriction to balanced trees, the cyclic permutation test cannot be used with most empirical datasets. However, because it is so conceptually straightforward and because it works (Fig. 2), it is a useful yardstick against which to measure another new approach in which permutations conserve the amount of phylogenetic signal in the data.

#### Signal-based permutations

The following permutation test can be used with real phylogenies: compare an empirical test statistic with the set of permutations in which phylogenetic signal is equal or sufficiently close to the empirical signal (Fig. 1C). The logic here is that the only rearrangements that can be meaningfully compared with empirical data are those in which trait values are just as conserved on, or structured by, the phylogeny. Phylogenetic signal is quantified here with Moran’s I rather than another metric like Pagel’s λ or Blomberg’s K in the non-parametric spirit of the permutation test: whereas those other metrics explicitly model the evolutionary process that generated a given trait, Moran’s I simply quantifies the degree to which the trait values of closely-related species covary (Gittleman and Kot 1990; Appendix 2). I implement signal-based permutation with a simple hill-climbing algorithm in which first the values of a trait are shuffled, then randomly-selected pairs of observations are swapped if doing so brings the phylogenetic signal closer to the empirical signal, and the procedure stops when the permuted phylogenetic signal is within some specified tolerance of the empirical value. The test could be implemented without this hill-climbing procedure, but it would make the test extraordinarily time-consuming for some datasets. Because phylogenetic signal depends on the values at the tips, the rearrangements that are included in the set of signal-based permutations depend on the values of the trait being permuted, unlike cyclic permutations and the LG permutation test. For applicable trees, the set of signal-based permutations is always at least as inclusive as the set of cyclic permutations: every cyclic permutation has identical phylogenetic signal, but non-cyclic permutations of a dataset can too (Fig 1B, rightmost permutation), and additional rearrangements can be accepted if the specified tolerance is large enough. For example, there are 2^(number of internal nodes) = 128 possible cyclic permutations of the 8-taxon tree in Figure 1B but 256 permutations with identical phylogenetic signal. The positions of clades (C,D) and (G,H) are switched in the 128 non-cyclic permutations.

Despite these striking differences from the cyclic permutation test, simulations show that signal-based and cyclic permutations have apparently identical statistical properties. Like the other test, the signal-based test correctly returns a uniform distribution of p values for 1000 simulations of uncorrelated evolution (Fig. 2, “Signal-based permutation”; Kolmogorov-Smirnov test against a uniform distribution, p = 0.413). The false negative rate is also comparable with that for the cyclic permutation test (Fig. 2; 716/1000 p-values below 0.05), and higher than that for PICs. Thus, the cyclic and signal-based permutation tests do not have the problems with statistical power and size that characterize the Lapointe-Garland test.

As a visual illustration of the two new phylogenetic permutation tests, consider their application to Felsenstein’s (1985) “worst case scenario” in which uncorrelated Brownian Motion evolution of two traits on a rooted tree of two polytomies with 20 tips each (all branch lengths equal) generates a spurious correlation among traits (Fig. 3A). An ordinary permutation test yields a distribution of mainly low absolute correlation coefficients (Fig. 3B) and a very high level of significance (p < 0.001). The investigator who makes the mistake of treating all tip values as exchangeable incorrectly rejects the null hypothesis of independent evolution. Conversely, the distribution of correlation coefficients for 1000 cyclic permutations is centered close to the empirical correlation coefficient (Fig. 3B), yielding a p-value of 0.31. Because of the clustering of trait values within subclades, every cyclic permutation preserves a relatively strong correlation coefficient: 95% of the cyclic permutations have |r| between 0.34 and 0.65. The null distribution generated by signal-based permutations depends on the tolerance: a set of 1000 signal-based permutations with the broadest possible tolerance (2) is statistically indistinguishable from an ordinary permutation test (Kolmogorov-Smirnov test, p = 0.7226) because the phylogenetic signal of every possible permutation is within its tolerance (Fig. 3B). For smaller tolerances, the distribution of test statistics for signal-based permutations more closely approximates the set of cyclic permutations (Fig. 3B), such that with a margin of 0.01 (Moran’s I of permuted variable Y between 0.512 and 0.532) they are statistically indistinguishable (p=0.536). Thus, signal-based permutations converge on the statistical properties of cyclic permutations.

**Figure 3.**
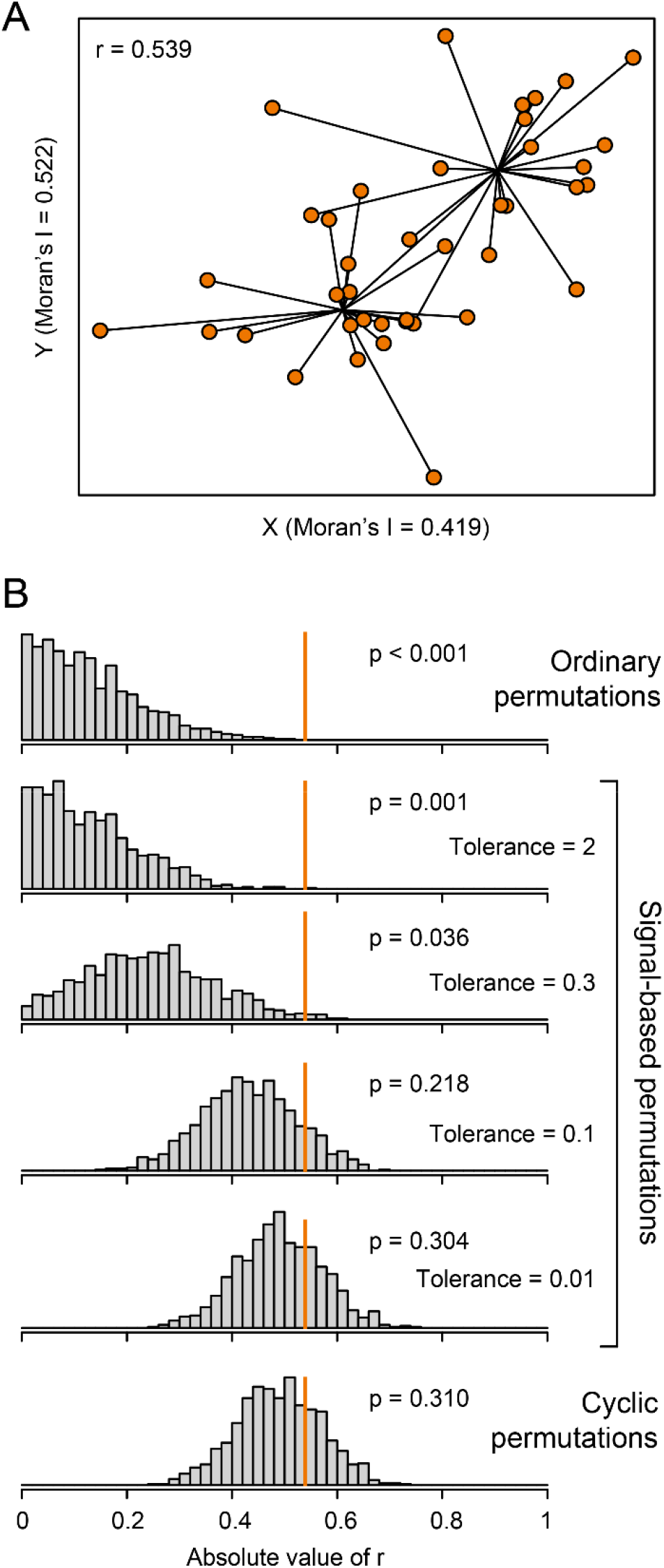
Phylogenetic permutations applied to Felsenstein’s “worst case” in which two traits evolve independently on a tree whose shape tends to induce spurious correlations. **3A.** Scatterplot of traits X and Y with lines connecting values at the tips to ancestral state reconstructions. **3B.** Histograms showing the correlation between X and Y for sets of 1000 permutations of the variable Y with different approaches: ordinary permutations, signal-based permutations with progressively smaller signal tolerances, and cyclic permutations. With a tolerance of 0.01, signal-based permutations are statistically indistinguishable from cyclic permutations. Vertical orange bars indicate the empirical correlation coefficient.

Interestingly, phylogenetic permutation tests succeed in a case where PICs and PGLS both fail: a second “worst case” constructed by Uyeda et al. (2018). In this scenario, simulated traits evolve in the same way and on the same phylogeny as in Felsenstein’s worst case, but with one modification: a single extreme shift in both traits near the root generates a contrast that is a strong enough outlier to make the two traits appear significantly associated, even when “correcting for phylogeny.” PICs/PGLS incorrectly recover significant relationships between traits because these methods are parametric, and their assumptions are violated by the dramatic outlier. The cyclic permutation test is unburdened by these assumptions: the extreme outlier is incorporated into every permutation, and the test correctly yields a non-significant result (Appendix 3). Likewise, the only rearrangements of the data that conserve phylogenetic signal are those in which exchanges only occur within clades and not between them, so a signal-based permutation test succeeds in the same way. Cyclic and signal-based permutations both represent reasonable null models against which to compare empirical patterns, but only the latter is applicable to real trees, so I use signal-based permutation tests to explore the following case studies.

### Case studies

The preceding sections established that phylogenetic permutations perform at least as favorably as PICs in “toy scenarios” in which the truth is known. These scenarios involved modeled BM evolution of traits with normal distributions, and the only test statistic considered was the correlation between two traits. PICs/PGLS perform comfortably within these bounds. In the following case studies, I use phylogenetic permutation to explore scientific questions that are effectively off-limits to PGLS-type methods because they involve strange distributions and test statistics beyond the least-squares regression framework. The first case study involves quantile regression on a heteroskedastic dataset with a non-normal response variable. The second explores the statistical significance of patterns of morphospace occupation. The third tests an ecogeographic rule: the data points are not tips in the phylogeny but the aggregate property of all the tips that occur in each geographic area.

The statistical significance of some of these findings could potentially be tested by comparing empirical statistics with null simulations rather than permutations, like what Mahler et al. (2013) used to demonstrate exceptional convergence in anoles. However, this requires assumptions about distributions and the evolutionary processes that generated a dataset which an investigator may not want or be able to make. If a dataset exhibits a more extreme test statistic than a set of simulations, is it because there was a mechanistic association between those traits, or because the simulations were unrealistic? Such questions may be hard to answer and are avoided by taking the non-parametric approach.

#### Quantile regression and peculiar distributions: arm number in feather stars

Saulsbury and Baumiller (2020) investigated a wedge-shaped relationship between absolute latitude and arm number among feather stars, a group of suspension-feeding marine echinoderms: species near the poles typically have around 10 arms, whereas those around the equator have between 5 and 150. Arm number varies widely within many families, but across the dataset it has a strange distribution, probably due to the unique and complex ontogeny of feather star arms (Shibata and Oji 2003): about half the species in the dataset have exactly 10 arms, and the rest of the distribution is markedly right skewed. More importantly, this non-normality also characterizes the residuals in a PGLS regression of arm number, and log(arm number), on absolute latitude. Another aspect of the dataset that poses obvious problems for least-squares regression is also the dataset’s most biologically interesting feature: arm number is heteroskedastic across absolute latitude. Beyond these more technical challenges, questions about the spread of a response variable as a function of a predictor cannot be readily addressed with least-squares regression. Instead they are the purview of quantile regression, which estimates quantiles (for example, the median, or the 10^th^ percentile) conditional on predictors. There is not currently an equivalent to quantile regression in the PGLS framework. As such, the authors used signal-based phylogenetic permutation tests to consider whether the latitudinal gradient in arm number could have plausibly emerged through independent evolution on feather star phylogeny.

Although both absolute latitude and arm number exhibit phylogenetic signal, and thus might be prone to spurious associations, the empirical relationships between the two are more extreme than almost all phylogenetic permutations. The 90^th^ and 95^th^ conditional percentiles, which characterize how maximum arm number relates to latitude, were significantly negative (p = 0.017 and 0.009, respectively), as was Spearman’s rank-correlation coefficient (p < 0.001). Concluding that the pattern could not be explained away as the result of random evolution, the authors drew on ecological and functional morphological evidence to argue that a latitudinal gradient in the intensity of predation represented the most plausible explanation for their findings. This simple case study illustrates the value of a comparative method that makes minimal assumptions about the distribution of the data. It also hints at the extent of the patterns that can be evaluated with phylogenetic permutation, although that is more fully illustrated by the following examples.

#### Morphospace occupation: Triassic ammonoids

Why are some theoretically possible morphologies not realized in nature, and why are some realized more frequently than others? These questions are the domain of theoretical morphology, a subdiscipline catapulted to the forefront of evolutionary biology for a time by David Raup. He found (1966) that the breadth of shell morphologies realized by mollusks and brachiopods was surprisingly well-summarized by a model in which a generating curve or whorl increases in size as it revolves around an axis. Shell geometry is controlled by three parameters: whorl expansion rate, translation of successive whorls along the axis, and the distance of successive whorls from the shell axis. Interestingly, most theoretically possible combinations of parameter values are not realized in nature; Raup cautiously submitted that either these unrealized forms were physiologically impossible, or shell-building invertebrates simply had not had time to reach those parts of morphospace yet. A companion paper (Raup 1967) focused on ammonoids, an extinct group of mostly “planispiral” mollusks in which typically no whorl translation occurs and variation is constrained along two axes of theoretical shell morphospace: the distance of successive whorls from the axis (D), and the whorl expansion rate (W) (Fig. 4A). Again, much of the rectangle defined by ammonoid occupation in D-W space is unoccupied – for example, almost no ammonoids fall above the line W = 1/D (Fig. 4A). Shells above this curve are open-coiled, making them, among other things, weaker and easier for a predator to crush. For shells under this curve, each whorl can incorporate part of the previous whorl in its construction, so open-coiled shells (W > 1/D) also waste the building materials they otherwise would have saved. Thus, the patterns in theoretical morphospace occupation are interesting because of the underlying fitness surface they suggest.

**Figure 4.**
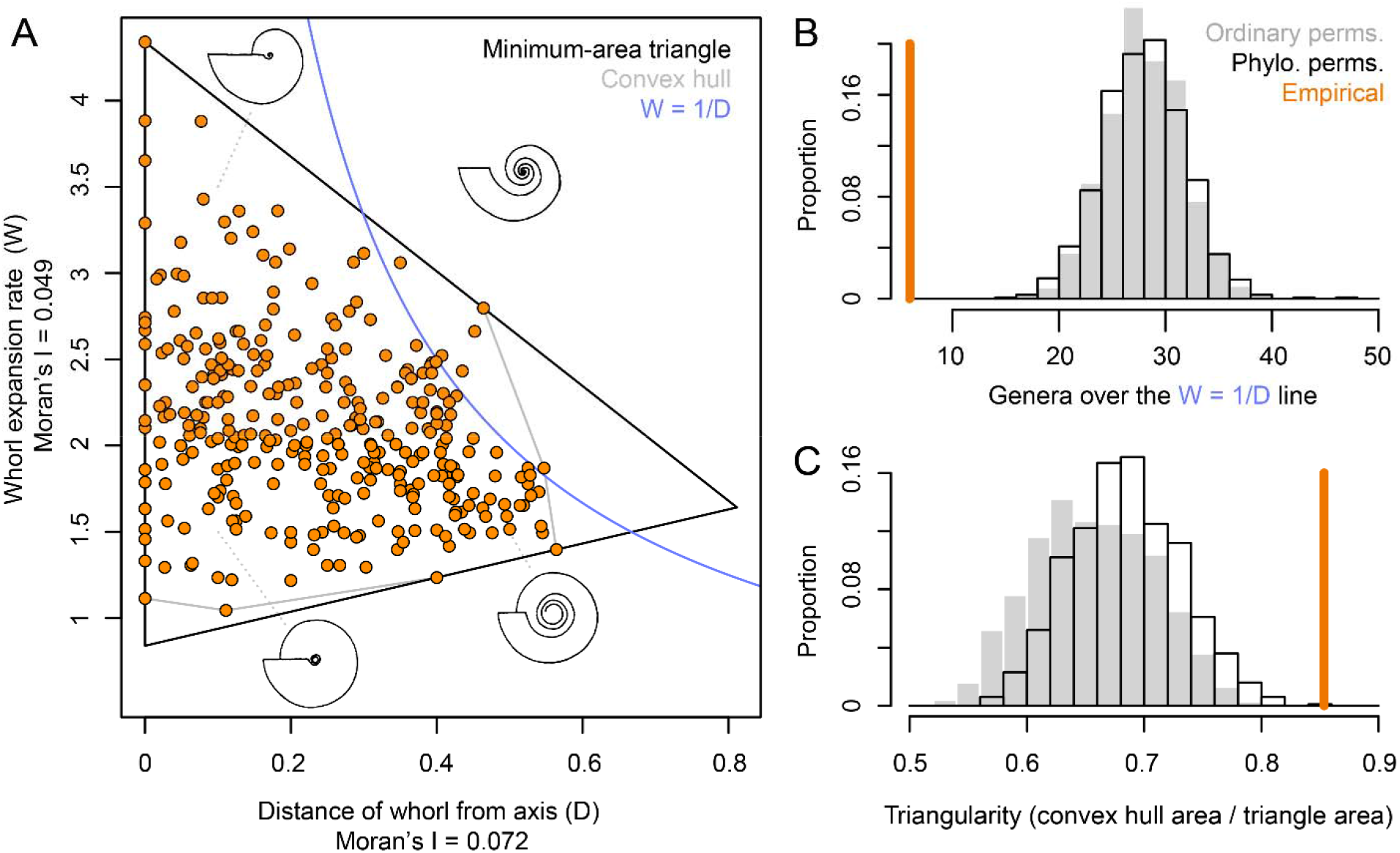
Phylogenetic permutation tests applied to theoretical morphospace occupation in Triassic ammonoids. **4A.** Two parameters controlling shell geometry in 322 Triassic ammonoid genera from McGowan (2004). Four theoretical ammonoid shells redrawn from Raup (1967) illustrate how different shell geometries correspond to different combinations of these parameters. Also plotted are the convex hull around the points, the smallest possible triangle around the points, and the line W = 1/D, above which shells are open-coiled. **4B.** The empirical number of genera with W > 1/D compared with the same statistic for 1000 ordinary and 1000 signal-based phylogenetic permutations. This statistic was taken by Raup (1967) as evidence for the reduced fitness of open-coiled forms. **4C.** The empirical ratio of the area of the convex hull around the data to the area of the smallest triangle that fits around the data, compared with the same statistic for 1000 ordinary and 1000 phylogenetic permutations. This metric of triangularity was interpreted by Tendler et al. (2015) in light of Pareto optimality theory.

The problem with inferences of selective forces from the pattern of morphospace occupation is that they rely on the equilibrium assumption (Lauder 1982): namely, that the phenotypes under study are at equilibrium with the selective forces that act on them. The alternate explanation for un- or under-occupied regions of morphospace is that, by chance, ammonoids simply have not had time to reach those regions yet – in other words, the system is historical and not at equilibrium. Raup (1967) raised this possibility, but admitted that in order to make headway he had to “assume that the observed morphology has had, in evolution, a selective advantage over other possible morphologies.” Subsequent studies have made the same assumption: for example, Tendler et al. (2015) tested whether ammonoids fill out a triangle in D-W space as a demonstration of Pareto optimality theory, which predicts that functional “archetypes” should form the vertices of polygons in trait space (Fig. 4A). They tested whether the ammonoid data are more triangular than the set of ordinary permutations, but this procedure incorrectly assumes that the data are exchangeable, or in other words that each data point obtained its morphology independently – a problem pointed out by Edelaar (2013) for another study of Pareto optimality. The equilibrium assumption leaves comparative studies vulnerable to the kinds of false positives discussed by Felsenstein (1985) in which a pattern apparently supported by a high number of replicates actually only represents a few evolutionary events. Theoretical morphology has not been incorporated with phylogeny in the way other comparative subdisciplines have in recent decades. However, it is not amenable to PGLS because it is not a regression problem: the question is not about the conditional mean of a response variable but about why certain combinations of traits are unrealized.

The phylogenetic permutation approach is a promising way forward for theoretical morphology because it can be used to ask what kinds of patterns in morphospace occupation can emerge without any dependence between traits. If empirical patterns fall outside the range of phylogenetic permutations, more interesting evolutionary explanations for the pattern in morphospace occupation can be explored – for example, certain morphologies could be unrealized because they are less fit. No broad-scale phylogeny of ammonoids is available, so I used taxonomy as a polytomy-rich phylogeny to explore morphospace occupation in the database of 322 Triassic ammonoid genera from McGowan (2004). These genera belong to 79 families in 18 superfamilies. This is a very coarse way to approximate phylogeny, so this exploration should be taken as a proof of concept and a hint at the role of contingency in ammonoid evolution.

I used ordinary and signal-based phylogenetic permutations to test the significance of two test statistics: the number of genera over the W=1/D line (6/322 genera), and the triangularity of the dataset in D-W space, defined as the ratio of the area of the convex hull to the area of the smallest triangle that encloses all the data (triangularity = 0.8535; Fig. 4A; Appendix 4). Both D and W have low signal on the “phylogeny”, with values of Moran’s I of 0.072 and 0.049, respectively. So, inasmuch as ammonoid taxonomy approximates phylogeny, the various ammonoid clades appear to have independently explored a lot of D-W space: for example, there are five superfamilies that each occupy more than half the area of the total convex hull. In a system characterized by this much exploration of morphospace, it seems unlikely that particularly strong patterns could emerge from random chance alone. The phylogenetic permutation test quantifies this preliminary impression: p < 0.001 for both test statistics for both ordinary and phylogenetic permutations (Fig. 4B-C). In other words, all phylogenetic permutations of the data have more open-coiled genera and are less triangular than the empirical dataset. Considering phylogeny (that is, going from ordinary to phylogenetic permutations) does not visibly affect the null distribution for the first test statistic; it does slightly for triangularity, shifting it to the right. So, because of phylogeny there is a slight tendency for permuted datasets to look more triangular, but not enough to make a difference for the p-value.

Thus, the independent evolution of shell growth parameters D and W constitutes a poor explanation for both the triangularity of the dataset and the paucity of open-coiled genera. One could easily imagine a hypothetical phylogenetic history for which a non-significant result would be obtained – for example, if every ammonoid with D > 0.3 were part of the same clade, it would be easier to explain away the pattern of morphospace occupation as a historical accident. The phylogenetic permutation test is well-suited for this problem because characterizing the biologically interesting features of morphospace occupation often requires the use of creative or novel statistics. Note that this is not the only way to evaluate the evolutionary “significance” of morphospace occupation: Tendler et al. (2015) showed that ammonoids refilled roughly the same region of morphospace several times after mass extinctions, representing semi-independent replicates. In the final case study, I explore a comparative problem in which the data points are not tips in the phylogeny.

#### Ecogeographic rules: Thorson’s rule in muricid gastropods

Some of the most productive hypotheses in biology predict the way some biological feature changes across space. Well-known examples of “ecogeographic rules” like these include Bergmann’s rule, the tendency for endotherms to be larger toward the poles (Olalla-Tárraga 2011), and Rapoport’s rule, the putative tendency for species’ latitudinal ranges to be smaller in the tropics (Stevens 1989). The analysis of ecogeographic rules entails an interesting and under-researched problem: species typically exist in more than one place, rather than at a single point as in other kinds of comparative studies. It might seem that a straightforward comparative analysis could address this by using a summary statistic of the range of each species, such as the range midpoint, and indeed many studies take this shortcut. However, such an approach removes biological information and is susceptible to false positives, especially if the trait in question corresponds with range size in some way (Saulsbury and Baumiller 2020; Colwell and Hurtt 1994). In the most well-known and straightforward example, a test for a relationship between absolute latitudinal midpoint and range size tends to recover strong negative relationships even none really exists: geometrically, large ranges cannot be centered at high latitude, so these taxa have their latitudinal midpoints “pulled” toward the equator (Colwell and Hurtt 1994). An alternative approach is to consider the ecogeographical data as such – that is, as a set of places and the aggregate properties of all the species in each place (Stevens 1989) – but this is analytically fraught as well. Such data are beset not only by the phylogenetic autocorrelation that complicates other comparative studies, but also by spatial autocorrelation to the degree that species occur in multiple places (Rohde et al. 1993). Here I show that both the phylogenetic permutation test can circumvent both sources of autocorrelation using a case study of larval development across latitude in muricid gastropods.

Thorson’s rule predicts that the larvae of marine invertebrates near the equator are more likely to be planktotrophs – feeding larvae that persist in the water column for a long time – whereas toward the poles there should be a predominance of non-feeding larvae, including direct developers and lecithotrophs (yolk-supplied larvae) (Thorson 1950). Thorson proposed that vulnerable planktotrophic larvae would not be able to cope with the extreme conditions and variable food supply at high latitudes, but this mechanism and the latitudinal pattern have subsequently received mixed empirical support (Marshall et al. 2012). Yet the idea persists: Pappalardo et al. (2014) claimed support for Thorson’s rule in a dataset of 44 muricid gastropod species (Fig 5A). A logistic PGLS regression of larval development (planktotrophic vs. non-feeding) on sea surface temperature [taken either from a single confirmed occurrence (69%) or from the latitudinal midpoint of each species (31%)] recovered marginally significant relationships: p = 0.087 for the regression of feeding (planktotrophic) vs. non-feeding mode on temperature, and p = 0.045 for the regression of pelagic (planktotrophic and lecithotrophic) vs. non-pelagic mode on temperature. Analyzing the same dataset, I found a similar degree of support in a PGLS logistic regression of larval development on latitudinal midpoints (Appendix 5). Other recent studies of Thorson’s rule use latitudinal or environmental midpoints as well (Ibáñez et al. 2018; Ewers-Saucedo and Pappalardo 2019), presumably because the PGLS framework requires it. Notably this seems to be a recent development, as Thorson and others who worked on this problem since were mostly considering the proportion of planktotrophic species at each latitude (Thorson 1950; Mileikovsky 1971; Jablonski and Lutz 1983; Collin 2003). Importantly, the use of midpoints can be vulnerable to complications involving range size: for example, if planktotrophic species have larger ranges, it would artificially strengthen the relationship between latitude and development by dragging the latitudinal midpoints of wide-ranging species toward the equator (Colwell and Hurtt 1994). In fact, the broad geographic ranges of feeding larvae are famous among invertebrate zoologists (Jablonski 1986), and the median latitudinal range of planktotrophic species in the muricid dataset is 3.25 times that of non-planktotrophs (Fig 5A). Using a latitude or temperature value selected randomly from the range might not be biased like the midpoint method is, but is not an ideal solution because it removes information and adds noise.

**Figure 5.**
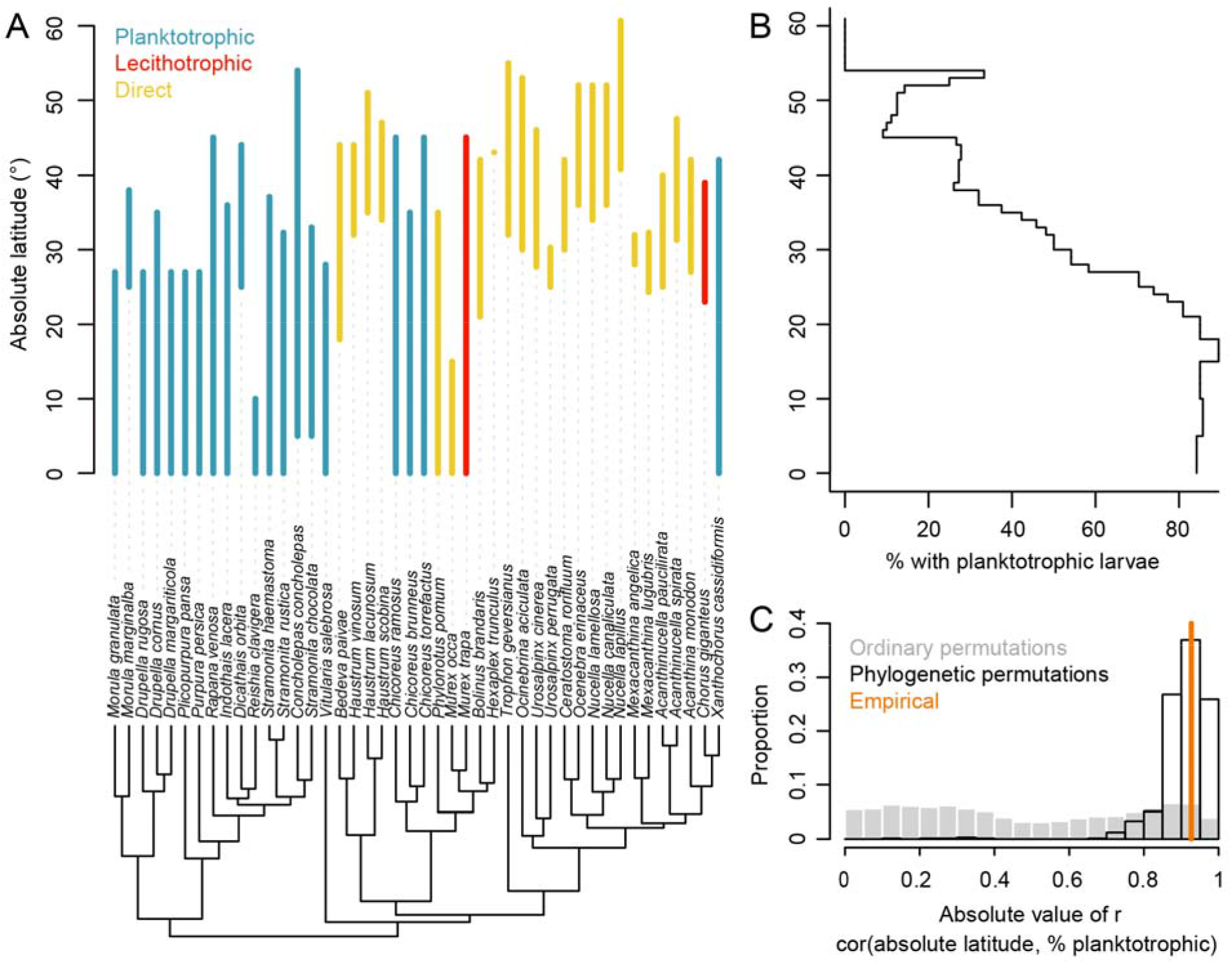
Phylogenetic permutation applied to Thorson’s rule in muricid gastropods. **5A.** Mode of larval development (color-coded), absolute latitudinal range, and phylogeny for the 44 species from Pappalardo et al. (2014). **5B.** Thorson’s rule plotted “as such”: the percentage of species with planktotrophic larval development in each 1° bin of absolute latitude. **5C.** The correlation between absolute latitude and the percentage of planktotrophic species in each 1° latitudinal bin, shown for the empirical data, 1000 ordinary permutations of mode of larval development, and 1000 phylogenetic permutations.

If instead the ecogeographic data are considered as such – for example, with a plot of the percentage of species with planktotrophic larvae in each 1° latitudinal bin – the trend is still apparent, with a strong correlation of r = 0.927 (Fig. 5B). A non-phylogenetic significance test that nevertheless accounts for spatial autocorrelation can be performed by permuting modes of larval development randomly across the tips of the phylogeny and re-computing the correlation coefficient (Fig. 5C). The resulting absolute correlation coefficients are spread evenly between 0 and 1, yielding marginal statistical significance with a p-value of 0.033. We can take both spatial and phylogenetic autocorrelation into account with signal-based phylogenetic permutations of mode of larval development. Phylogenetic signal of planktotrophic vs. non-planktotrophic development is high (Moran’s I = 0.60), and indeed larval development only appears to have transitioned on the phylogeny a few times, providing an investigator with very low sample size: the most parsimonious history of development involves only three transitions to or away from planktotrophy. Accordingly, almost all phylogenetic permutations have high absolute correlation coefficients, from which the empirical correlation is statistically indistinguishable (p = 0.387). Thus, the muricid dataset cannot provide strong evidence against the completely independent evolution of latitude and larval development. Notably, the authors focused on temperature not latitude; it is unclear if a similarly non-significant result would be obtained for the correlation between temperature and larval development, but temperature and latitude are closely correlated, and the same analytical problem applies because species occupy a range of temperatures.

Ecogeographic data present an interesting challenge to the comparative biologist because the data points, cast most directly, do not represent tips in the phylogeny but the aggregate properties of all the tips in the phylogeny that occur in each place. It might be possible to consider such data in a PGLS framework, but it would require the specification of a rather complex variance-covariance matrix. Crucially, this phylogenetic permutation test does not provide evidence *against* Thorson’s rule in this group. At the pattern level, the group is a clear example of the rule, with a strong negative correlation between latitude and the proportion of species with planktotrophic larvae. This trend probably has important implications for their modern ecology and future evolution, because it predicts, for example, that low-latitude planktotrophic species should be buffered against extinction by their broad ranges (Jablonski and Lutz 1983; Jablonski 1986). However, the key point is that, with phylogenetic and spatial autocorrelation this strong, such a trend could have easily arisen without any mechanistic relationship between latitude and larval development. In fact, given the distribution of phylogenetic permutations (Fig. 5C), it would be much more surprising to find no latitudinal trend in larval development. This might explain why so many groups appear to follow the rule (Ibáñez et al. 2018), especially since mode of larval development appears to evolve infrequently among marine invertebrates (Collin 2004). It would require an exceptionally strong trend to support a mechanistic Thorson’s rule in a dataset like this one – or more plausibly, a different kind of data. This might mean a clade in which larval development transitions more frequently, or it might mean a different kind and scale of evolutionary repetition.

## Conclusions

The regression-based approach to comparative biology has been hugely successful, but it is also inflexible: it fails for strangely distributed response variables, but more importantly, the range of questions it can address is limited. Permutation tests represent a powerful alternative that performs well both within and outside the bounds where PGLS is applicable. Case studies illustrate the use of phylogenetic permutations for pushing comparative methods to new places.

Rather than being a purely technical matter, the distinction between PGLS and permutation-based approaches is underlain by a more substantive difference in attitude toward comparative data. PGLS is typically described as a way to “correct for phylogeny” (Symonds and Blomberg 2014). Other comparative methods take an even more direct approach by transforming the data to “remove phylogenetic effects” (Stearns 1984; Cheverud et al. 1985; Felsenstein 1985; Gittleman and Kot 1990). The implication is that comparative data have been contaminated or affected by an agent called phylogeny, and that this contamination needs to be isolated and removed before the real relationships in the data can be studied. It is a drawback of these methods that they put the user at a remove from the raw data. Patterns in phylogenetically autocorrelated data are also no less real than those in transformed data: biological phenomena that could have arisen purely by chance, like Thorson’s rule in some taxa, can nevertheless have real and important consequences. Transformations and corrections also remove information and limit the kinds of statistics and questions that can be applied to a dataset.

The phylogenetic permutation test is mostly unique among comparative methods in that it treats the raw data as such. The test is subject to some of the same criticisms to which all frequentist tests are subject, including that statistical significance tells an investigator nothing about effect size (a reaction to the widespread conflation of “significance” with importance; Dushoff et al. 2019). This is true, but for many biological phenomena including the case studies discussed here, the most relevant effect size is arguably the empirical test statistic. Only six of 322 Triassic ammonoid genera have open-coiled shells; it is hard to imagine a more meaningful phylogenetic transformation of this test statistic. The phylogenetic permutation framework, which considers whether raw data look typical for cases of independent evolution, is in a way the reverse of the reigning paradigm of transforming comparative data or their expected covariances to fit into a regression analysis. Hopefully, these new approaches can help facilitate scientific creativity among comparative biologists.

## Acknowledgements

This paper benefited greatly from critical reviews from Tomasz K. Baumiller, Kelly K.S. Matsunaga, Caroline Parins-Fukuchi, James B. Pease, and Nathanael Walker-Hale. I thank Avichai Tendler and Paula Pappalardo for providing data from their studies on ammonoids and muricid gastropods, respectively.

## Data availability statement

Code, supplementary data, and appendices are available at the following GitHub repository: https://github.com/jgsaulsbury/phyloperm

