## Appendix for "Permutation tests for comparative data"

**Appendix 1: Lapointe-Garland permutations**

A high rate of auto-exchanges in Lapointe-Garland (LG) permutations has the effect of up-weighting rearrangements in which observations are not moved from their original position. This causes test statistics applied to Lapointe-Garland (LG) permutations to be more tightly constrained around the empirical test statistic than the other two phylogenetic permutation tests, which should reduce statistical power. As an illustration, consider LG permutations performed on a rooted, balanced, dichotomizing tree of four taxa A-D. This illustration will consider the most phylogenetically informed implementation of LG permutations in which k=1. LG permutations exchange the values at two tips with probability proportional to the phylogenetic distance between those tips; taxa A and B are separated from C and D by the greatest distance possible in the tree, so no exchanges will be made between them. Thus, LG permutations will generate four arrangements: ABCD, BACD, ABDC, and BADC. Each taxon has a 1/3 probability of being exchanged with its sister, and a 2/3 chance of staying put. Thus, the probability of an LG permutation yielding the first of the above four rearrangements is 2/3 * 2/3 = 4/9; for the middle two, it is 2/3 * 1/3 = 2/9; and for the last, it is 1/3 * 1/3 = 1/9. Simulations bear this out: the observed proportions in 100,000 simulations are 0.445. 0.223, 0.221, and 0.110. Notably, cyclic and signal-based permutations do not have this differential weighting feature.

This weighting feature has the effect of pulling permuted test statistics closer to the empirical one relative to the other kinds of phylogenetic permutation. Take for example the following dataset:

|  | A | B | C | D |
| --- | --- | --- | --- | --- |
| X | 1 | 2 | 3 | 4 |
| Y | 1 | 2 | 3 | 4 |

The empirical correlation between X and Y is 1. Cyclic and signal-based permutations weight all rearrangements equally, yielding a permuted absolute correlation of 0.6 for one quarter of permutations, 0.8 for half of the permutations, and 1 for the last quarter. LG permutations of Y yield correlations closer to the empirical one: r = 0.6 for the just 1/9 of permutations, r = 0.8 for 4/9, and r = 1 for the last 4/9. This extremely simplified example shows why test statistics are “dragged” toward the observed statistic by LG permutations. The same effect should be observed in empirical datasets. The reduction of statistical power that comes with this effect can be observed in Fig. 2 in the main text.

**Appendix 2: Moran’s I**

Moran’s I is a metric of autocorrelation, originally devised to measure spatial autocorrelation (Moran 1950) but outfitted by Gittleman & Kot (1990) for quantifying the degree to which the trait values of closely-related species covary. Moran’s I sums the weighted averages of each observation’s covariation with every other observation; weights quantify relatedness among observations. At least one study has taken the weight for the covariance between tips *i* and *j* (*w_i,j_*) to be inverse of the patristic distance between the two (ex. Münkemüller et al. 2012). With this scheme, the weight for the covariation between tips *i* and *j* initially decays rapidly with increasing patristic distance, then decays more slowly for larger distances. It is not clear why the Moran’s I metric needs to exaggerate the distances between closely related species and dampen distances between distantly related ones. Thus, this study sets *w_i,j_* equal to the amount of shared phylogenetic history between *i* and *j* as a proportion of the largest root-to-tip distance in the tree.

**Appendix 3: Uyeda’s worst case**

Uyeda et al. (2018) considered an modification of Felsenstein’s famous “worst case” in which two simulated traits evolve in the same way on the same tree, but a single extreme shift in both traits near the root generates a contrast that is strong enough to make the two traits appear to be associated with phylogenetically independent contrasts (PICs). I compare the performance of PICs with that of cyclic and signal-based permutations using the exact settings as used in this paper for Felsenstein’s worst case, except that a shift value is added to all the members of one subclade for each of the two traits. This shift value is drawn from a normal distribution with mean zero and variance equal to 1000 times the rate parameter for the simulated traits, following Uyeda et al. (2018). Also, the tree is a balanced 8-taxon tree rather than a 40-taxon tree for the sake of speed.

Correlations between PICs for the two traits were erroneously found to be statistically significant (p < 0.05) in 990 out of 1000 simulations generated following the protocol above. Conversely, p values for cyclic and signal-based permutations appear uniformly distributed, with 51/1000 and 46/1000 tests returning p < 0.05, respectively. Phylogenetic permutations thus represent a meaningful null against which to compare evolutionary phenomena with strange features that violate the assumptions of parametric PCMs.

**Appendix 4: Case study, Triassic ammonoids**

In this paper, we calculate the “triangularity” of the data as the ratio of the area of the convex hull enclosing a bivariate dataset to the area of the smallest triangle that fits around that dataset. The smallest triangle will always be at least as big as or bigger than the convex hull; they will obviously be identical if the convex hull is a triangle. Shoval et al. (2013) quantified triangularity in this way in their study of Pareto optimality across various groups. In a subsequent study focusing on ammonoids (Tendler et al. 2015), the same research group used a different metric of triangularity involving archetypal analysis. The two approaches return similar results, and the former is used in this study because it is more straightforward. The smallest triangles enclosing sets of data points were calculated with the *minboundtri* function in the MATLAB package “MinBoundSuite” (D’Errico 2020).

**Appendix 5: Case study, Thorson’s rule in muricid gastropods**

Pappalardo et al. (2014) used logistic PGLS to recover marginal statistical support for a relationship between temperature and mode of larval development in muricid gastropods. Analyzing the same dataset with the same technique, I recovered statistically significant relationships between absolute latitude (midpoint or randomly selected; see main text) and feeding vs. non-feeding larval development (b_1_ estimate = -0.058, p = 0.017) and pelagic vs non-pelagic development (b_1_ estimate = -0.086, p = 0.0020).
